# The significance of social interactions in synchronized swarming flight in a termite

**DOI:** 10.1101/2023.12.25.573318

**Authors:** Nobuaki Mizumoto, Tomonari Nozaki

**Affiliations:** Okinawa Institute of Science & Technology Graduate University, Onna-son, Okinawa, Japan; Laboratory of Insect Ecology, Graduate School of Agriculture, Kyoto University, Kyoto, Japan; Laboratory of Evolutionary Genomics, National Institute for Basic Biology, Okazaki, Aichi, Japan

**Keywords:** Animal collective behavior, Self-organization, Synchronization, Temporal patterns

## Abstract

In social insect colonies, individuals of working caste coordinate their actions to manage various collective tasks. Such collective behaviors are observed not only in working castes but also in reproductives. During certain seasons, newly emerged winged reproductives (alates) fly from the nest to disperse and find mating partners in a synchronized manner. Although this “swarming” behavior is one of the collective behaviors that involve the greatest number of individuals in social insects, underlying social interactions remain unexplored. Here, we show that synchronized flight among colony members results from collective decision-making. Our simple simulation model suggests that social interactions in a large group of alates enable synchronized flight within a nest under fluctuating environmental conditions. This model is supported by empirical experiments using a termite *Reticulitermes kanmonensis*. Under the semi-field environment with fluctuating temperatures, alates within the same colony synchronized their dispersal flight under higher air temperatures, while dispersal flight was suppressed under lower temperatures. Furthermore, termites could synchronize their dispersal flights even under laboratory conditions with constant temperature, indicating that environmental cues are not always necessary for synchronization. In either case, higher synchronization happened with a larger number of alates. All these results demonstrate that both environmental and social factors interplay to enable the synchronized swarming flight of social insects.

## Introduction

Social insects present a variety of coordinated collective behavior and accomplish social tasks involving many individuals. Such ability to solve group-level problems is called swarm intelligence (Garnier et al., 2007), and many modeling and empirical studies revealed the individual-level behavioral mechanisms underlying social phenomena (Camazine et al., 2001). In insect society, most such collective behavior is performed by working castes, such as workers and soldiers. On the other hand, it may not be widely recognized, but the collective behaviors of social insects are not limited to social contexts. During mating, many ant and termite species convergently evolved “swarming flight,” where numerous newly emerged alates (winged individuals) come out of the nests, fly to disperse, and obtain mating partners in a highly synchronized manner (Nutting, 1969). Although subsequent mating behavior is entirely different between ants and termites, synchronized swarming flight could be adaptive for both taxa by reducing predation due to predator satiation (Moser et al., 2004; Nutting, 1979), increase the likelihood of mate finding (Hölldobler and Wilson, 1990), and promote outbreeding between colonies (Aguilera-Olivares et al., 2015; Husseneder et al., 2006; Noordijk et al., 2008). Although the role of social interactions in one-time behavioral synchronization is widely studied in various group-living animals (e.g., synchronized hatching (Endo et al., 2019)), social interactions underlying the swarming flight behavior remain poorly understood.

Both environmental cues and behavioral coordination within a group should be critical for synchronized swarming flight. Previous studies have identified the environmental cues that elicit a swarming flight by social insects (Hölldobler and Wilson, 1990; Nutting, 1969). In termites, for example, high temperature is a good predicting factor of the occurrence of swarming flights in some species (Chouvenc et al., 2017; Puckett et al., 2014; Sugio et al., 2020), while rainfall and humidity are essential triggers for species especially in arid (Aihetasham and Akhtar, 2008) or tropical regions (Lucena et al., 2022; Mitchell, 2008; Neoh and Lee, 2009). The environmental cues are sometimes complex, where air temperature and relative humidity are not the only factors (Martius, 2003; Sands, 1965). In any case, such environmental cues are shared among different colonies in the same area, thus essential for synchronization among colonies. On the other hand, within a colony, groups of alates need to make a consensus to synchronize flight timing, and social interactions must play an important role. Without interactions, synchronization within a colony is equivalent to that among colonies; if all colony members respond to the environmental cues independently, synchronization cannot happen under a constant environment.

A termite *Reticulitermes kanmonensis* Takematsu 1999 is an ideal model for the experimental operation of swarming behavior. As in most other termite species, alates of this species fly to disperse from their original nests as a group during a limited time of the year (Takematsu, 1999). This species, however, is unique regarding the time lag between the emergence of alates and flight dispersal. Unlike other species, whose alates disperse as soon as they emerge, alates of this species emerge in November, wait within a nest over the winter, and fly to disperse from late February to April next coming year (see Table S1 for observational data in this study, and Table S2 for the summary of all literature information), similar to *R. flaviceps* (Khan et al., 2019). During the waiting period, alates are ready for mating behavior and colony foundation (Supplementary Text, Fig. S1). Thus, by collecting colonies during winter and incubating them in various laboratory conditions, we can study the factors affecting synchronization of mating swarming.

In this study, we investigate the role of social interactions in synchronizing the swarming flight among colony members. First, we developed a simple mathematical model to predict how social interactions can suppress small flight events and enhance synchronized large swarming events. Second, by observing the swarming dynamics of *R. kanmonensis* under uncontrolled semi-natural environments and controlled experimental conditions, we demonstrate that *R. kanmonensis* uses temperature to determine the swarming date but can synchronize swarming even without that. Finally, by exploring the relationship between colony size and swarming synchronicity, we propose that the termite swarming flight is a self-organized collective behavior regulated by social interactions.

### Model prediction

In social insect colonies, all individuals are not identical but usually have variability in their threshold to respond to the same environmental cues (e.g., (Ishii and Hasegawa, 2013)). This variability must also be applied to alates because the alates within a nest are heterogenous physiological states due to endogenous variability, e.g., sexes, timing of the final molt, nutritional status, and genotypes. Suppose each alate in the nest responds to the current temperature to determine if it disperses on that day. As each alate has a different responsive threshold, some alates should become active for dispersal, while others should remain inactive at a certain temperature (Fig. 1A). Therefore, to achieve synchronized swarming flight, alates within a nest must achieve consensus.

**Figure 1.**
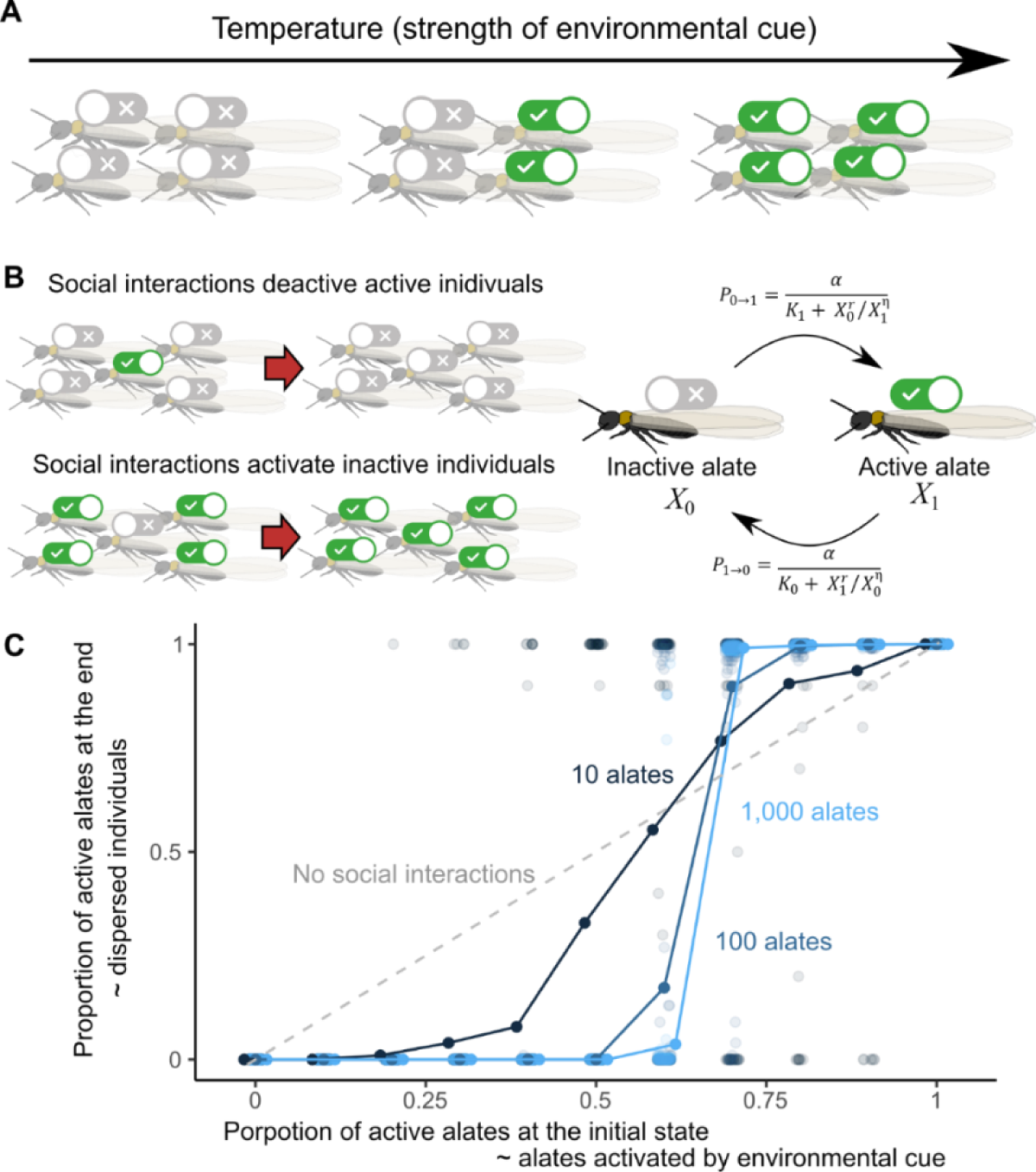
A framework of collective decision-making model applied to termite swarming flight behavior. (A) An assumption that a group of alates within a nest have different response thresholds. At an intermediate temperature, some individuals are activated while others are not. (B) A simple stochastic model for synchronized alate activation. Active individuals surrounded by inactive individuals are deactivated, while inactive individuals surrounded by active individuals are activated. (C) The results of the simulations. For each parameter (group size *n* and proportion of active alates at the initial state), we ran 100 simulations. The parameters were set as `α` = 0.01, *K*_0_ = 2, *K*_1_ = 1, and `η` = 2.

To study this process, we employed a simple collective decision-making model of social insects (Jeanson et al., 2012) and extended it to swarming flight behavior. In this model, each alate becomes active/inactive states based on the environmental cues (i.e., temperature) (Fig. 1A) and then decide if they change their states based on the social interactions (Fig. 1B). The probability *P*_1→0_ of changing their state from active (state 1) to inactive (state 0) as a function of the number of active individuals (*X*_0_) and inactive individuals (*X*_1_) is defined by:

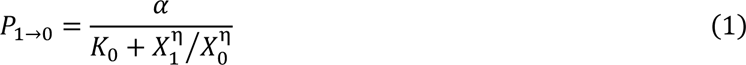

and the probability *P*_0→1_ of changing their state from inactive (state 0) to active (state 1) is:

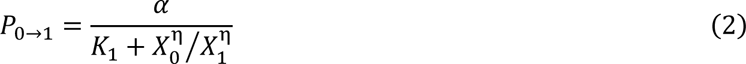

where constant `α` controls the speed of state changes, constant *K*_0_ and *K*_1_ represent the intrinsic attractiveness of each state: the higher the value of *K*, the more termites prefer to be in that state, and the parameter `η` controls the steepness of the response to other alates, i.e., the degree of non-linearity: the higher the value of `η`, the greater the influence of other alates on the individual decision to change their states.

In the Monte Carlo simulations, the proportion *r* of total *n* alates initially becomes active for swarming. Then, each alate decides if they change their state based on the equation (1) or (2). After this process was iterated 1,000 times, we counted the number of active individuals (who will fly to disperse from the colony) and inactive (who will remain in the nest). We investigate variable proportion *r* (0, 0.1, 0.2, 0.3, 0.4, 0.5, 0.6, 0.7, 0.8, 0.9, 1.0) and total alates number *n* (10, 100, 1,000). For each parameter set, we ran 100 simulations to record the number of alates that decided to disperse. The simulation program was implemented in R (R Core Team, 2023).

As a result, our model demonstrated that social interactions enhance flight synchronicity (Fig. 1C). Without interactions, only alates that are activated by environmental cues disperse from the nest (y=x, grey dashed line in Fig. 1C). In this scenario, synchronization can happen when the super strong environmental cue exists; however, the weak environmental cue can result in minor dispersal events. Social interactions can suppress these minor dispersal flight events (Fig. 1C) and enable them to synchronize the timing of swarming with enough strong environmental cues. The effect of social interactions is stronger with a larger number of alates.

## Material and Methods

### Termite collection

The species *R. kanmonensis* was described from Japan (Takematsu, 1999) but originated in southern China via anthropogenic introductions to Japan (Kitade and Hayashi, 2002). It is also native to Taiwan (C.-C. Wu et al., 2019) and was introduced to Korea from Japan (Lee et al., 2015). We collected colonies of *R. kanmonensis* with a piece of nesting wood in Ube city and Sanyo-Onoda city in Yamaguchi prefecture, Japan. The field collections were performed twice, 7-8th December 2016 and 15th-16th February 2017. This region has both *R. kanmonensis* and *R. speratus* (Kitade and Hayashi, 2002). Among the 42 colonies we observed, all 12 colonies with alates were *R. kanmonensis*, while all 30 colonies without alates but with last-instar nymph were *R. speratus*. The list of the colonies is in Table S1.

### Semi-field swarming experiment

To observe the natural swarming flight of *R. kanmonensis* colonies, we observed termite colonies under a daily fluctuating environment, using six colonies collected in February (I-VI). For each colony, we placed the pieces of wood nested by termites in a plastic container without a lid. Then, we placed this container inside a larger plastic container (Fig. 2A). This apparatus enabled alates to come out of the smaller plastic container during swarming flight and prevented them from returning to the original nests simultaneously. We installed this observational container in a PVC greenhouse on the North Campus of Kyoto University. We observed if any alates came out of the nest once a day after sunset (∼6 pm) for 40 days. We examined the mating behavior and colony foundation success of these alates (Supplementary Text, Fig. S1). At the end of all observations, we opened the nests using knives and Japanese hatchets to confirm that no alates remained in the nest.

**Figure 2.**
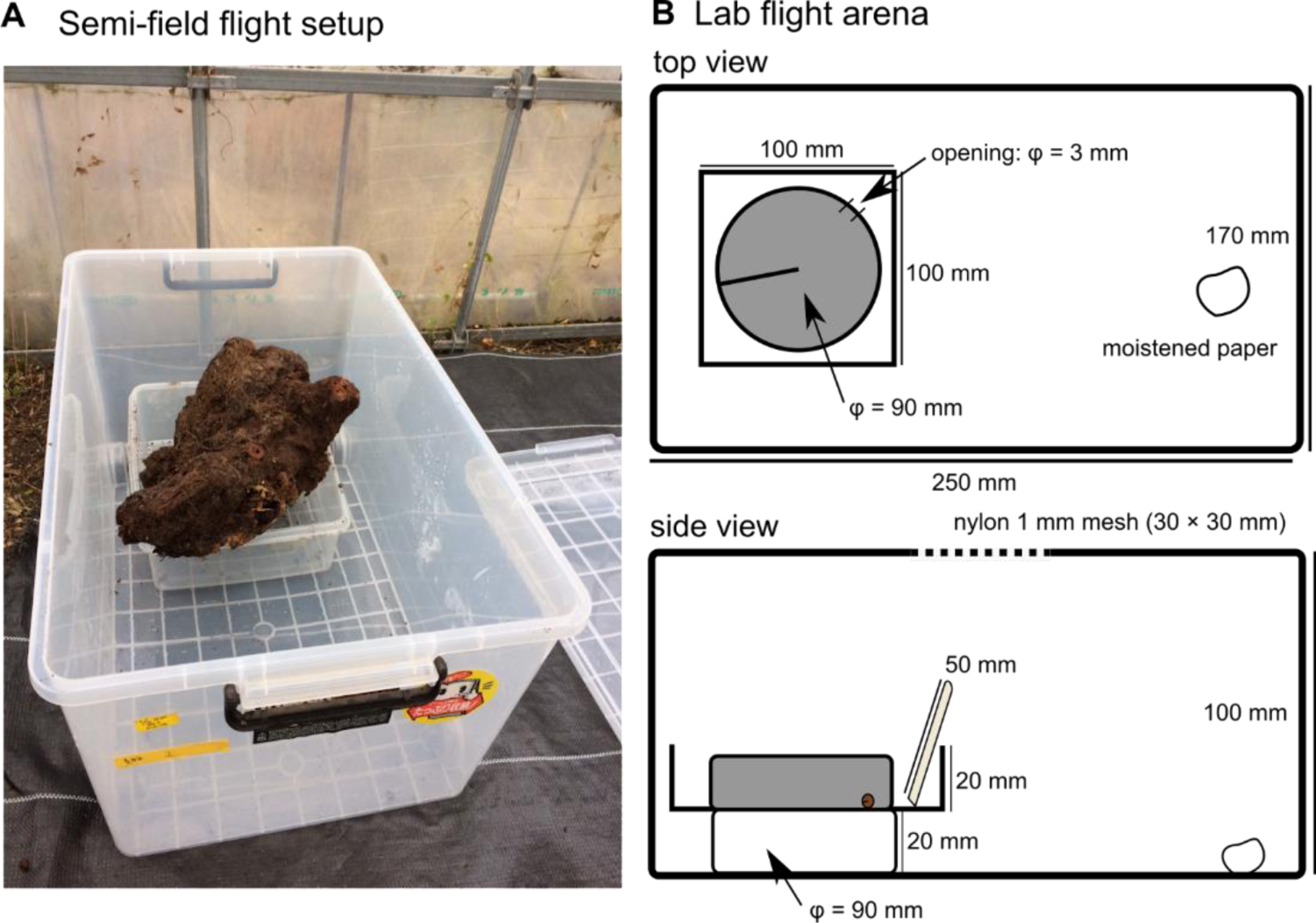
Experimental setups for termite swarming observations. (A) A photo of the setup for the semi-field experiment. Termite nesting wood is located in a plastic container without a lid, which is installed inside a larger plastic container. (B) Drawings of the setup for laboratory observations.

### Laboratory swarming experiment

We investigated the termite swarming flight under constant temperature conditions, using five colonies (A-E) collected in December. Soon after returning from the field (within two days), we carefully opened the nesting wood with knives and Japanese hatchets and separated all termite individuals, including alates, larvae, soldiers, and workers. Then, we measured the weight of these individuals (with contaminated small wood chips) and prepared subgroups that had a weight ranging from 10g to 20g. We obtained 12 different subgroups (two for A, B, C, and E; four for D). All these processes were performed in a cooled room (10℃ ± 2℃) to minimize the disturbance for termite alates. We put each subgroup in a Petri dish (∮ = 90mm) filled with brown-rotted pinewood mixed cellulose medium (Mitaka et al., 2023) up to 5mm in depth. We wrapped the dish with aluminum foil to keep the inside dark. This makes termites nest within the dish. Each dish has a hole (∮ = 3 mm) on the side so that termite alates can come out of the nest for dispersal flight (Fig. 2B). We installed the dish and disposal wooded chopstick (cut 50mm in length) in a plastic container (100×100×20mm) without a lid. This apparatus enables only alates to come out of the container while workers and soldiers stay inside. Furthermore, fixing this apparatus on another 90 mm dish (20mm in height) prevented dispersed alates from returning to the original nest. Then, we placed this in a plastic container (170×250×100mm) for observation. We kept each subgroup under a constant light environment (12L12D) and a constant temperature. We prepared two different temperature treatments (5-10℃ or 20℃); the former simulated the winter environment, while the other simulated the spring environment. We checked if alates came out of the apparatus once a day at the end of the light period as this species swarms around noon in the field (Takematsu, 1999). If any alates came out of the apparatus, we collected them, separated them by sex, and recorded the number. In case workers came out of the apparatus accidentally, we gave them back to the nest. We examined the mating behavior and colony foundation success of these alates (Supplementary Text) and confirmed that these alates are equivalent to those under semi-natural conditions (Fig. S1). The observation lasted for 70 days. At the end of all observations, we opened the artificial dish nest and confirmed that no alates remained.

### Statistical analysis

First, we determined a major dispersal flight event (= swarming flight). In a dataset of the number of alates that came out of the nest, we identified outlines using the z-score method. We calculated the z-score of the number of alates using the *scale()* function in R, and then we identified the data with z-score > 3 as outliers (Figs. 3A, 4A). Then, we compared the proportion of alates dispersed on the swarming day between semi-field and laboratory conditions as a degree of synchronicity of swarming. Also, we counted the number of dispersed alates retroactively to estimate the number of alates within a nest for each observational date. Using this estimation, we investigated the effect of the total number of alates present on the degree of synchronicity. We used a generalized linear mixed-effect model (GLMM) with binomial errors and a logit-link function, where swarming conditions (lab or semi-filed) and the total number of alates were treated as fixed effects, while the original colony was included as a random effect (random intercept). To test the statistical significance of the inclusion of each explanatory variable, we used the likelihood ratio test (type II test).

**Figure 3.**
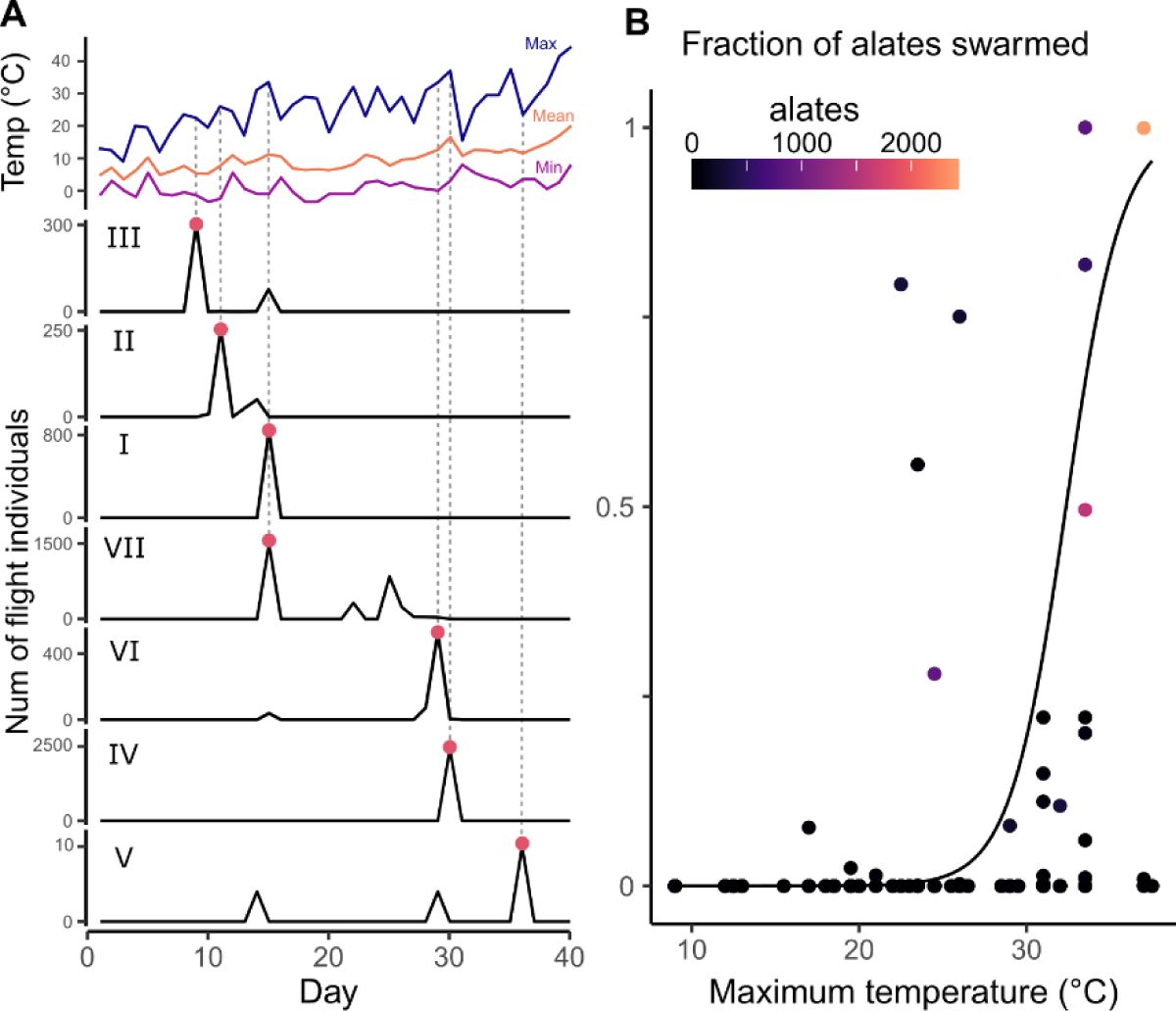
Swarming flight in semi-natural conditions. (A) Time development of the temperature within the greenhouse and the number of alates dispersed from the nest for each colony. Red points indicate the swarming (= major dispersal) events (z-score > 3). (B) Relationship between maximum temperature and the fraction of alates dispersed from the nest. The solid line is the regression curve calculated with GLMM, indicating a significant positive relationship (*P* < 0.001).

**Figure 4.**
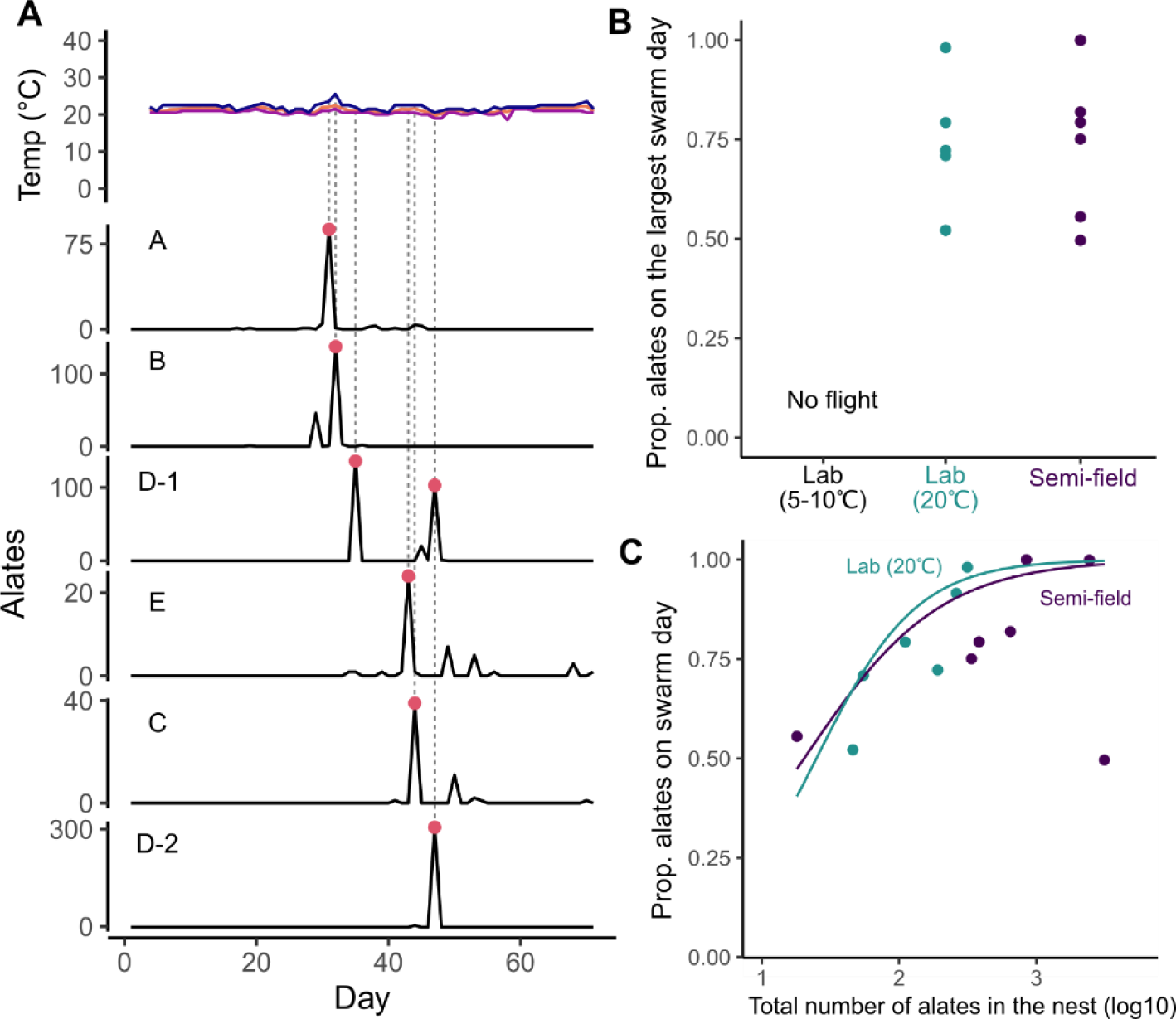
Swarming flight in laboratory conditions. (A) Time development of the temperature within the incubator and the number of alates dispersed from the nest for each colony. Red points indicate the swarming (= major dispersal) events (z-score > 3). (B) Comparison of the proportion of alates dispersed on the largest dispersal date. (C) Relationship between the total number of alates in the nest and the proportion of alates dispersed during swarming events.

For semi-field observations, we investigated the relationship between daily temperature and the number of dispersing alates using a GLMM with binomial errors and a logit-link function. For each day, we calculated the number of alates present inside the colony. Then we included the binomial decision of each alate to disperse (1) or remain in the nest (0) in the model. The average, maximum, or minimum temperature of each day was treated as a fixed effect, and the original colony was included as a random effect (random intercept).

All statistical analyses were conducted using R v4.3.1 (R Core Team, 2023) with the *glmer()* function in the package “lme4” (https://cran.r-project.org/web/packages/lme4/index.html) and likelihood ratio test with the *Anova()* function in package “car.” All data and R code are available at GitHub (github.com/nobuaki-mzmt/Ret-kanmonensis-swarming). The accepted version will be uploaded to Zenode to obtain DOI for the version of record.

## Results and Discussion

Under the semi-natural conditions with variable temperatures across days, all nests had just one swarming event (Fig. 2A). The occurrence of dispersal alate depended on the temperature. More alates dispersed from the nest on the day with higher maximum temperature (GLMM; `χ`^2^_1_ = 9363.1, *P* < 0.001; Fig. 2B), while swarming is suppressed on the day with lower temperature. We obtained the same results using average or minimum temperature (GLMM; *P* < 0.001), but the AIC was smallest with maximum temperature. The observed relationship between the highest temperature and swarming events is consistent with field observations in other termite species (e.g., (Chouvenc et al., 2017; Puckett et al., 2014; Sugio et al., 2020)). Two of the seven colonies observed had the swarming event on the same day (colonies I and VII). Also, four colonies swarmed within two-day differences (Colonies II and III, Colonies IV and VI), which is close enough to form a pair (Kusaka and Matsuura, 2017). Thus, termites synchronized swarming timing between colonies, using temperature cues.

Even under laboratory conditions with constant daily temperature, colonies of *R. kanmonensis* showed within colony synchronization and swarming flight (Fig. 4A). When the room temperature was kept as low as the winter air temperature (5-10℃), termites did not show any dispersal flights (Fig. 4B). However, when the temperature was kept at as high as the spring swarming season (20℃), all alates dispersed from the original nests. Even though the temperature was constant in our experiments, all six observed groups showed at least one synchronized swarming event (Fig. 4A), and the proportion of individuals dispersed on the largest swarm day was not different from semi-field conditions with variable temperature (Fig. 4B). Therefore, for within-colony synchronization of swarming flight, environmental cue of temperature is not always necessary. Instead, social interactions among colony members should have played a crucial role, as predicted by our model (Fig. 1). Note that no synchronization between colonies was observed in the laboratory condition. Thus, environmental cues are essential for across-colony synchronization.

In self-organized collective behavior, group size is an important parameter regulating group-level dynamics (Dornhaus et al., 2012). Our model predicted that within-colony synchronization of swarming flights is facilitated by larger colony size (Fig. 1). This is consistent with our observations under both semi-natural and laboratory conditions (Fig. 4C). In our experiments, the total number of alates in the nest was highly variable across colonies, where the strength of synchronization (the proportion of alates that dispersed on the largest swarming date) had the positive relationship with the total number of alates that existed on the date of swarming (GLMM, alate number: `χ`^2^_1_ = 5.98, *P* = 0.014, experimental conditions: `χ`^2^_1_ = 0.08, *P* = 0.78, interaction: `χ`^2^_1_ = 0.11, *P* = 0.74, Fig. 4C). Interestingly, this relationship was present in both semi-natural conditions and laboratory conditions with constant temperature (Fig. 4C). This strongly indicates that within-colony synchronization of dispersal flight is mediated by social interactions in *R. kanmonensis*; with large colonies, termite alates can synchronize dispersal flight even without environmental cues.

Our finding on the association between colony size and the synchronicity of swarming has a vital implication for understanding the evolution of termite mating behavior (Figs 1 and 4). The mating behavior of termites is diverse across lineages (Mizumoto et al., 2022a; Nutting, 1979). Some species show a short and concentrated swarming season in dispersal flight, where dispersal flights are highly synchronized within and across colonies (Nutting, 1979). On the other hand, some species show a long swarming season with less synchronized dispersal flights distributed over several months. The latter is often the case in one-piece nesting termites (e.g., (Sugio et al., 2020)), whose colony life is completed within a piece of wood that serves as food and nest (Abe, 1987; Mizumoto and Bourguignon, 2020). Phylogenetic comparative analysis showed that one-piece nesting termites have smaller colony sizes than other nesting types (Mizumoto et al., 2022b), and thus, their total number of alates within a nest must be smaller. Because synchronization is not facilitated by social interactions when the number of alates is small (Fig. 1), species with smaller colony sizes may not use synchronized swarming flights as a mating mechanism but develop other unknown alternative behavioral mechanisms. Interestingly, with several exceptions, tandem running behavior is lacking for several one-piece nesting termite genera, such as, *Bifiditermes*, *Neotermes*, *Pterotermes* and *Stolotermes* (summarized in (Mizumoto et al., 2022a)).

As a player, alates should be the most important agents for synchronized swarming events— however, actual swarming results from more complex social interactions involving different castes within a nest. For example, in advance of swarming events, workers excavate tunnels towards the surface of the wood, soil, or mound nests and build structures from which alates fly to disperse (Nutting, 1969) (Fig. 5). Occasionally, chambers for waiting alates can be built by workers near the entrance, especially in mound-building termites (e.g., (Sen-Sarma, 1962)). The proportion of the soldiers in the nest reaches the maximum before the swarming season (Haverty and Howard, 1981), where soldiers usually are associated with alates outside of the nests at the time of swarming, presumably defending colony members and alates from predators (Fig. 5; Fig. 1 of (Mizumoto and Bourguignon, 2022)). Similarly, worker-alates interactions also play an important role in swarming among ants (Hölldobler and Wilson, 1990). The research on swarming flight from social behavior perspectives remains descriptive works, and further studies are required to understand how colonies achieve consensus decision-making regarding the timing of swarming flight.

**Figure 5.**
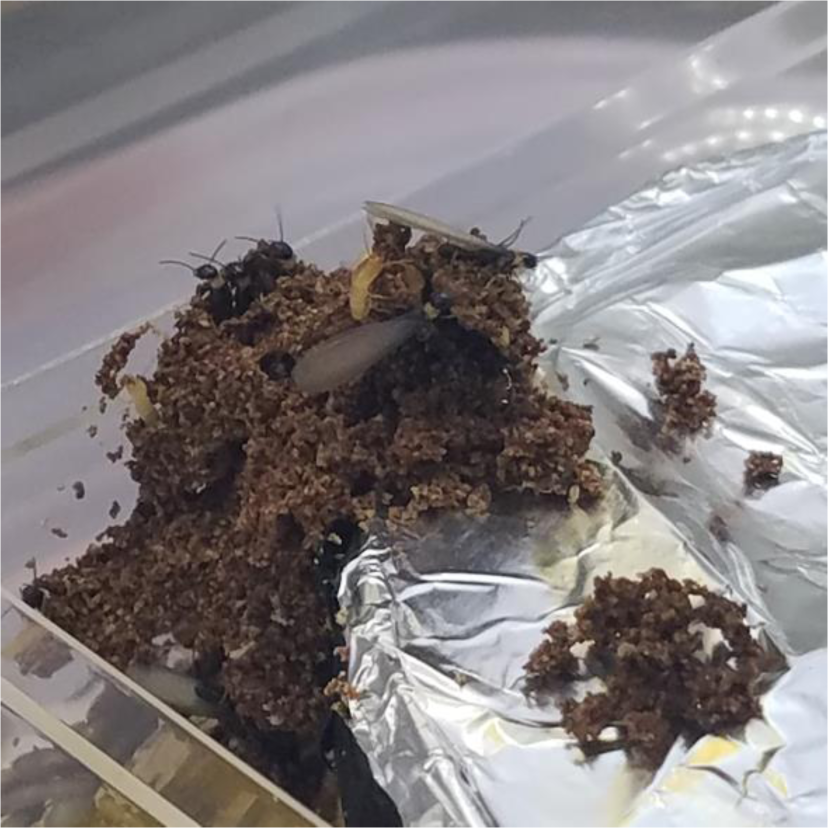
Preflight behavior in *R. kanmonensis*. Termites build shelter tubes from the experimental apparatus in Fig. 2B as openings for swarming. Several alates with workers and soldiers came out to sense the external environment.

Our study highlights the possibility of experimentally manipulating termite swarming behaviors. Alates of *R kanmonensis* are ready to mate during winter (Supplementary Text, Fig. S1), but they do not start dispersal flight under low-temperature conditions (5-10℃, Fig. 4B). Thus, by changing temperatures, the timing of swarming can be easily manipulated, making this species the ideal model to study termite mating biology. For example, *R. kanmonensis* was introduced to Japan and Korea and sympatric with *R. speratus* (Kitade and Hayashi, 2002; Lee et al., 2015). As alates of *R. kanmonensis* overwinter after emergence and disperse earlier in the year than *R. speratus*, these two species are separated in the timing of flight seasons, avoiding hybridization between them. In this study, we could test if these two species have the physiological potential to produce hybrid colonies by delaying swarming flights of *R. kanmonensis* until alates of *R. speratus* become ready for mating (Supplementary Text, Fig. S2). This is consistent with other hybrid combinations in *Reticulitermes* termites (e.g., (J. Wu et al., 2019; Wu et al., 2023)).

In conclusion, here we study the synchronized swarming flight of a termite as a collective decision-making system to suggest that activation/deactivation feedback between alates plays a central role in achieving synchronization. In social insects, a similar self-organized mechanism is used in different contexts of collective decision-making by a group of workers, including aggregation, foraging, and nest site selection (Jeanson et al., 2012). Furthermore, our model can be easily extended to different one-time synchronized behavioral events that involve inter-individual communications in other taxa, such as synchronous hatching by insects (Endo et al., 2019; Nishide and Tanaka, 2016), birds (Vince, 1966) and crocodiles (Vergne and Mathevon, 2008). Thus, this study contributed to understanding the diversity of animal collective behavior by adding the overlooked example of the reproductive castes of social insects. Synchronized swarming flight is the convergent collective behavior in social insects, especially ants and termites. Although the success of swarming flight is critical for their life history, few studies have experimentally manipulated this behavior to understand its ultimate and proximate causes. Further comparative studies, especially between different degrees of synchronization across species, will reveal the evolutionary origin of the swarming flights, which had affected the evolution of food web by providing the excessive number of preys in a short period.

## Acknowledgments

We appreciate Kenji Matsuura at Kyoto University for valuable suggestions, and Kaoru Watanabe for experimental assistance. This study was supported by the JSPS Research Fellowships for Young Scientists DC1 (15J02767 to N.M., and 16J08955 to T.N.). Also, modeling was supported by a Grant-In-Aid for Early-Career Scientists from JSPS (21K15168). N.M. was supported by an IPSF fellowship from OIST.

## Author Contributions

N.M.: conceptualization, data curation, formal analysis, funding acquisition, investigation, methodology, project administration, resources, supervision, validation, visualization, writing-original draft, writing-review and editing. T.N.: conceptualization, funding acquisition, investigation, methodology, writing-review and editing.

## Competing Interest Statement

The authors declare that they have no conflicts of interest in the contents of this manuscript.

## The File includes

-Table S1. *Reticulitermes* colonies collected from Yamaguchi prefecture

-Table S2. Records of dispersal flights in *R. kanmonensis*.

-Supplementary Text

-Figure S1. Comparison of colony foundation success between different wintering conditions.

-Figure S2. Comparison of colony foundation success in heterospecific pairs.

**Table S1.** *Reticulitermes* colonies collected from Yamaguchi prefecture.

| Date | Species | Location | Latitude | Longitude | Composition |
| --- | --- | --- | --- | --- | --- |
| Dec. 7, 2016 | *R. kanmonensis* | Fujioyama, Sayama, Yamaguchi | 34.0328 | 131.3842 | Alate |
| Dec. 7, 2016 | *R. kanmonensis* | Fujioyama, Sayama, Yamaguchi | 34.0323 | 131.3829 | Alate |
| Dec. 7, 2016 | *R. kanmonensis* | Ryuoan, Sanyo-Onoda, Yamaguchi | 33.9512 | 131.1725 | Alate |
| Dec. 8, 2016 | *R. kanmonensis* | Ejio, Sanyo-Onoda, Yamaguchi | 34.0435 | 131.1871 | Alate |
| Dec. 8, 2016 | *R. kanmonensis* | Eiandai, Sanyo-Onoda, Yamaguchi | 34.0149 | 131.1303 | Alate |
| Dec. 7, 2016 | *R. speratus* | Fujioyama, Sayama, Yamaguchi | 34.0321 | 131.3838 | Nymph |
| Dec. 7, 2016 | *R. speratus* | Aiofutajima, Yamaguchi, Yamaguchi | 34.0373 | 131.4048 | Nymph |
| Dec. 7, 2016 | *R. speratus* | Ryuoan, Sanyo-Onoda, Yamaguchi | 33.9509 | 131.1700 | Nymph |
| Dec. 7, 2016 | *R. speratus* | Ryuoan, Sanyo-Onoda, Yamaguchi | 33.9512 | 131.1698 | Nymph |
| Dec. 7, 2016 | *R. speratus* | Ryuoan, Sanyo-Onoda, Yamaguchi | 33.9510 | 131.1725 | Nymph |
| Dec. 7, 2016 | *R. speratus* | Ryuoan, Sanyo-Onoda, Yamaguchi | 33.9569 | 131.1681 | Nymph |
| Dec. 7, 2016 | *R. speratus* | Ryuoan, Sanyo-Onoda, Yamaguchi | 33.9510 | 131.1698 | Nymph |
| Dec. 8, 2016 | *R. speratus* | Ejio, Sanyo-Onoda, Yamaguchi | 34.0363 | 131.1895 | Nymph |
| Dec. 8, 2016 | *R. speratus* | Ejio, Sanyo-Onoda, Yamaguchi | 34.0365 | 131.1893 | Nymph |
| Dec. 8, 2016 | *R. speratus* | Ejio, Sanyo-Onoda, Yamaguchi | 34.0371 | 131.1922 | Nymph |
| Dec. 8, 2016 | *R. speratus* | Ejio, Sanyo-Onoda, Yamaguchi | 34.0378 | 131.1952 | Nymph |
| Dec. 8, 2016 | *R. speratus* | Ejio, Sanyo-Onoda, Yamaguchi | 34.0427 | 131.1867 | Nymph |
| Dec. 8, 2016 | *R. speratus* | Ejio, Sanyo-Onoda, Yamaguchi | 34.0416 | 131.1854 | Nymph |
| Dec. 8, 2016 | *R. speratus* | Ejio, Sanyo-Onoda, Yamaguchi | 34.0413 | 131.1856 | Nymph |
| Dec. 8, 2016 | *R. speratus* | Ejio, Sanyo-Onoda, Yamaguchi | 34.0388 | 131.1855 | Nymph |
| Dec. 8, 2016 | *R. speratus* | Ejio, Sanyo-Onoda, Yamaguchi | 34.0332 | 131.1872 | Nymph |
| Dec. 8, 2016 | *R. speratus* | Ejio, Sanyo-Onoda, Yamaguchi | 34.0330 | 131.1872 | Nymph |
| Dec. 8, 2016 | *R. speratus* | Ejio, Sanyo-Onoda, Yamaguchi | 34.0364 | 131.1892 | Nymph |
| Dec. 8, 2016 | *R. speratus* | Ejio, Sanyo-Onoda, Yamaguchi | 34.0369 | 131.1891 | Nymph |
| Dec. 8, 2016 | *R. speratus* | Eiandai, Sanyo-Onoda, Yamaguchi | 34.0153 | 131.1299 | Nymph |
| Dec. 8, 2016 | *R. speratus* | Shimofuriyama, Ube, Yamaguchi | 34.0115 | 131.2668 | Nymph |
| Dec. 8, 2016 | *R. speratus* | Shimofuriyama, Ube, Yamaguchi | 34.0119 | 131.2672 | Nymph |
| Dec. 8, 2016 | *R. speratus* | Ajisu, Ube, Yamaguchi | 34.0212 | 131.3208 | Nymph |
| Date | Species | Location | Latitude | Longitude | Composition |
| Feb. 15, 2017 | *R. kanmonensis* | Ryuoan, Sanyo-Onoda, Yamaguchi | 33.9518 | 131.1732 | Alate |
| Feb. 15, 2017 | *R. kanmonensis* | Ryuoan, Sanyo-Onoda, Yamaguchi | 33.9514 | 131.1725 | Alate |
| Feb. 15, 2017 | *R. kanmonensis* | Ryuoan, Sanyo-Onoda, Yamaguchi | 33.9515 | 131.1724 | Alate |
| Feb. 15, 2017 | *R. kanmonensis* | Ryuoan, Sanyo-Onoda, Yamaguchi | 33.9516 | 131.1731 | Alate |
| Feb. 15, 2017 | *R. kanmonensis* | Hamagochi-Ryokuti, Sanyo-Onoda, Yamaguchi | 33.9459 | 131.1803 | Alate |
| Feb. 16, 2017 | *R. kanmonensis* | Motoyamamisaki, Sanyo-Onoda, Yamaguchi | 33.9315 | 131.1810 | Alate |
| Feb. 16, 2017 | *R. kanmonensis* | Iwaya-no-hana, Yamaguchi, Yamaguchi | 33.9867 | 131.3957 | Alate |
| Feb. 15, 2017 | *R. speratus* | Ryuoan, Sanyo-Onoda, Yamaguchi | 34.0326 | 131.3839 | Nymph |
| Feb. 15, 2017 | *R. speratus* | Hamagochi-Ryokuti, Sanyo-Onoda, Yamaguchi | 33.9437 | 131.1789 | Nymph |
| Feb. 15, 2017 | *R. speratus* | Hamagochi-Ryokuti, Sanyo-Onoda, Yamaguchi | 33.9436 | 131.1789 | Nymph |
| Feb. 15, 2017 | *R. speratus* | Hamagochi-Ryokuti, Sanyo-Onoda, Yamaguchi | 33.9454 | 131.1803 | Nymph |
| Feb. 15, 2017 | *R. speratus* | Hamagochi-Ryokuti, Sanyo-Onoda, Yamaguchi | 33.9521 | 131.1833 | Nymph |
| Feb. 16, 2017 | *R. speratus* | Ejio, Sanyo-Onoda, Yamaguchi | 34.0353 | 131.1867 | Nymph |
| Feb. 16, 2017 | *R. speratus* | Ejio, Sanyo-Onoda, Yamaguchi | 34.0354 | 131.1867 | Nymph |

**Table S2.** Records of dispersal flights in *R. kanmonensis*. Almost all information was summarized in (1, 2).

| Event | Condition | Year | Date | Place | Reference |
| --- | --- | --- | --- | --- | --- |
| Emergence | field | 1911 | 11/21 | Chofu, Yamaguchi | (3) |
|  | field | 1912 | 10/27-11/3 | Chofu, Yamaguchi | (4) |
|  | field | 1912 | 10/25 | Shimonoseki, Yamaguchi | (4) |
|  | field | 1916 | 10/28 | Shimonoseki, Yamaguchi | (5) |
| Dispersal flight | field | 1911 | 3/26-3/31 | Moji, Fukuoka | (6) |
|  | field | 1911 | 2/28-3/1 | Kokura, Fukuoka | (6) |
|  | field | 1911 | 3/18 | Moji, Fukuoka | (7) |
|  | Lab | 1914 | 2/16 |  | (8) |
|  | Lab | 1915 | 3/6 |  | (8) |
|  | Lab | 1916 | 1/19 |  | (8) |
|  | Lab | 1917 | 3/13 |  | (8) |
|  | Lab | 1919 | 2/26 |  | (8) |
|  | Lab | 1919 | 3/19 |  | (8) |
|  | Lab | 1920 | 3/9 |  | (8) |
|  | Lab | 1920 | 3/23 |  | (8) |
|  | Lab | 1921 | 4/1, 4/4 |  | (9) |
|  | Lab | 1922 | 2/23, 2/25 |  | (10) |
|  | Lab | 1922 | 4/1-4/4 |  | (11) |
|  | Lab | 1923 | 3/15, 20 |  | (12) |
|  | unknown | 1925 | 4/2 |  | (13) |
|  | field | 1928 | 3/31 | unknown | (14) |
|  | field | 1911 | 12/24 | Taiwan | (7) |
|  | field | 1911 | Mid-Nov | Taiwan | (7) |

Nawa used a pen name “Konchu-okina (昆虫翁)” in several papers in Konchu Sekai (昆虫世界).
Journal name Konchu Sekai (昆虫世界) is also called “The insect world.” Here we used Konchu Sekai.
We also list original titles because several reports published in the same year by the same author have very similar titles, which makes it challenging to locate the correct references only with translated titles.

1. S. Ito, “Kanmon-shiroari” (1). *Shiroari* **57**, 3–10 (1984). 伊藤修四郎, 関門白蟻〔1〕. しろあり 57, 3–10 (1984).
2. S. Ito, “Kanmon-shiroari” (2). *Shiroari* **58**, 2–8 (1984). 伊藤修四郎, 関門白蟻〔2〕. しろあり **58**, 2–8 (1984).
3. U. Nawa, Kyushu region termite survey team, returns. *Konchu Sekai* **16**, 20 (1912). 名和梅吉, 再び九州地方白蟻調査団. 昆蟲世界 **16**, 20 (1912)
4. U. Nawa, Naptual flight of termite that emerges earlier. *Konchu Sekai* **16**, 455 (1912). 名和梅吉, 羽化の早き白蟻. 昆蟲世界 **16**, 455 (1912)
5. U. Nawa, Emergence season of *Reticulitermes kanmonensis*. *Konchu Sekai* **20**, 468–469 (1916). 名和梅吉, 関門白蟻の羽化期. 昆蟲世界 **20**, 468–469 (1916)
6. U. Nawa, On “kiashi-shiroari” (*Reticulitermes flaviceps*). *Konchu Sekai* **15**, 194–195 (1911). 名和梅吉, 黄肢白蟻に就て. 昆蟲世界 **15**, 194–195 (1911)
7. U. Nawa, On a termite that emerges earlier. *Konchu Sekai* **16**, 33 (1912). 名和梅吉, 羽化の早き白蟻に就いて. 昆蟲世界 **16**, 33 (1912).
8. U. Nawa, Comparison of swarming periods of “Kanmon-Shiroari” (*Reticulitermes kanmonensis*). *Konchu Sekai* **24**, 137 (1920). 名和梅吉, 関門白蟻群飛時期の比較. 昆蟲世界 **24**, 137 (1920)
9. U. Nawa, Swarming flight of “Kanmon-Shiroari” (*Reticulitermes kanmonensis*). *Konchu Sekai* **24**, 137 (1921). 名和梅吉, 関門白蟻の群飛. 昆蟲世界 **24**, 137 (1921)
10. U. Nawa, Swarming flight of “Kanmon-Shiroari” (*Reticulitermes kanmonensis*). *Konchu Sekai* **26**, 92–93 (1922). 名和梅吉,関門白蟻の群飛. 昆蟲世界 **26**, 92–93 (1922)
11. U. Nawa, Another swarming flight of “Kanmon-Shiroari” (*Reticulitermes kanmonensis*). *Konchu Sekai* **26**, 129–130 (1922). 名和梅吉, 再び関門白蟻の群飛. 昆蟲世界 **26**, 129–130 (1922)
12. U. Nawa, Another swarming flight of “Kanmon-Shiroari” (*Reticulitermes kanmonensis*). *Konchu Sekai* **27**, 135 (1923). 名和梅吉, 再び関門白蟻の群飛. 昆蟲世界 **27**, 135 (1923)
13. U. Nawa, Greeting from mosquito, silkworm, and termite alates. *Konchu Sekai* **29**, 134–135 (1925). 名和梅吉, 蚊蚕と羽蟻のお見舞. 昆蟲世界 **29**, 134–135 (1925)
14. U. Nawa, Flight of “Kanmon-Shiroari” (*Reticulitermes kanmonensis*). *Konchu Sekai* **31**, 131 (1928). 名和梅吉, 関門白蟻の飛行. 昆蟲世界 **31**, 131 (1928)

## Supplementary Text

### Mating behavior of wintering alates in *R. kanmonensis*

#### Methods

We investigated if alates dispersed under the laboratory condition (Fig. 2B) function as well as alates dispersed under the semi-natural conditions. In *Reticulitermes* termites, dispersed alates shed their wings and search for mating partners. Once a female and a male are encountered, they perform tandem runs, with the male following the female to look for a potential nest site. We investigated if females and males form tandem running pairs across different conditions. We placed a female and a male, which shed their wings by themselves, on a petri dish filled with moistened plaster (`φ` = 90 mm). We recorded their behavior for four minutes and checked if pairs engaged in tandem runs for more than 30 seconds. The behavioral observations were performed for 69 pairs from six colonies in total. All pairs were from the same colony, and each individual was observed only once. Pairing colony mates does not affect the tandem running success in *R. speratus* (Mizumoto et al., 2022a).

Furthermore, we investigated the colony foundation success of alates across three different conditions: (i) just after collected during the winter (18 pairs in total), (ii) after dispersal flight in laboratory conditions (71 pairs in total), and (iii) after dispersal flight in semi-natural conditions (73 pairs in total). Pairs of a female and a male were chosen randomly and placed on a block of brown-rotted pinewood mixed cellulose medium (Mitaka et al., 2023) (25mm diameter × 10mm height) at the center of a petri dish (diameter = 65 mm). All pairs were created using nestmate individuals. Petri dishes were maintained at 25℃ in the dark for 60 days. After 60 days, we disassembled the sawdust block and counted the surviving individuals for each caste. We defined the colony with both a male and a female surviving as succeeding in the colony foundation.

To analyze the success of colony foundation, we used GLMM with binomial errors and a logit-link function. The condition of individuals (wintering, lab flight, semi-field flight) was treated as a fixed effect, and the original colony was included as a random effect (random intercept). Next, to investigate whether the different conditions of individuals would affect the number of produced offspring using GLMM with Poisson errors. For this analysis, we omitted the data of colonies that failed in colony foundation. Similarly, the condition of individuals (wintering, lab flight, semi-field flight) was treated as a fixed effect, and the original colony was included as a random effect (random intercept). To test the statistical significance of the inclusion of each explanatory variable, we used the likelihood ratio test (type II test). In cases of significant effects of the time, we ran Tukey’s post hoc test. All GLMM analyses were conducted using R v4.3.1 (R Core Team, 2023) with the *glmer()* function in the package “lme4”, likelihood ratio test with the *Anova()* function in the package “car” and post hoc test with the *glht()* function in the package “multcomp”.

#### Results

Tandem running behavior was observed in all 69 pairs that swarmed in the laboratory condition, indicating that the experimental setup in Fig. 2B could successfully simulate the swarming in *R. kanmonensis*. We found a significant difference in the proportion of colony foundation success among conditions of individuals (GLMM, LRT, `χ`^2^ = 8.22, *P* = 0.016, Fig. S1). The colony success was higher in lab-swarming individuals than semi-natural swarming individuals (TukesHSD, *Z* = −2.66 *P* = 0.02). On the other hand, once termites succeeded in colony foundation, there were no significant differences in the number of offspring among conditions of individuals (GLMM, LRT, `χ`^2^_2_ = 5.24, *P* = 0.073, Fig. S1).

**Figure S1.**
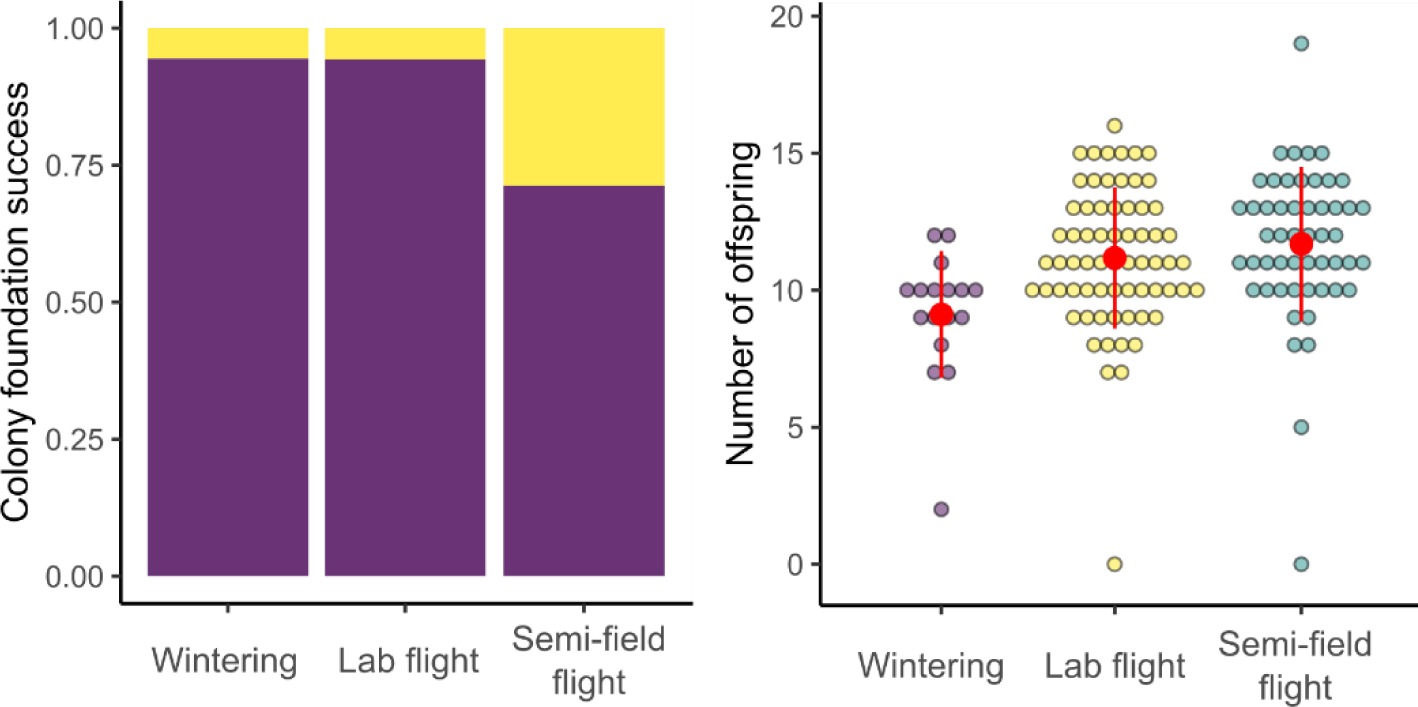
Comparison of colony foundation success between different wintering conditions.

### Hybridization between *R. kanmonensis* and *speratus*

In the studied area, *R. kanmonensis* is the introduced species and sympatric with *R. speratus*. As alates of *R. kanmonensis* overwinter after emergence and disperse earlier in the year than *R. speratus*, these two species are separated in the timing of flight seasons, avoiding hybridization between them. However, these two species have the physiological potential to produce hybrid colonies. In this study, we found that alates of *R. kanmonensis* do not initiate dispersal flight under 5-10℃ conditions (Fig. 4B), and thus, the swarming of *R. kanmonensis* could be delayed until alates of *R. speratus* become ready for mating. Using this feature, we compared tandem running behavior, colony foundation success, and the number of offspring between conspecific pairs and hybrid pairs. We used one colony of *R. kanmonensis* and *R. speratus* to make a pair of Rkf (*R. kanmonensis* female)-Rkm (x10), Rkf-Rsm (x9), Rsf-Rkm (x10), and Rsf-Rsm (x10). We evaluated their tandem running behavior and colony foundation success with the same methods as above.

All hybrid combinations could form tandem running pairs. We found that both hybrid combinations could successfully establish colonies and produce offspring. Especially, there was no significant difference in the colony foundation success among pairing combinations (GLMM, LRT, `χ`^2^_3_ = 3.37, *P* = 0.338, Fig. S2). On the other hand, after colony foundation success, we found a significant difference in the number of offspring (GLMM, LRT, `χ`^2^_3_ = 9.21, *P* = 0.027, Fig. S2), where Rkf-Rsm had a bit smaller number of offspring than Rkf-Rkm pairing (Tukey HSD, *Z* = 0.168, *P* = 0.045).

**Figure S2.**
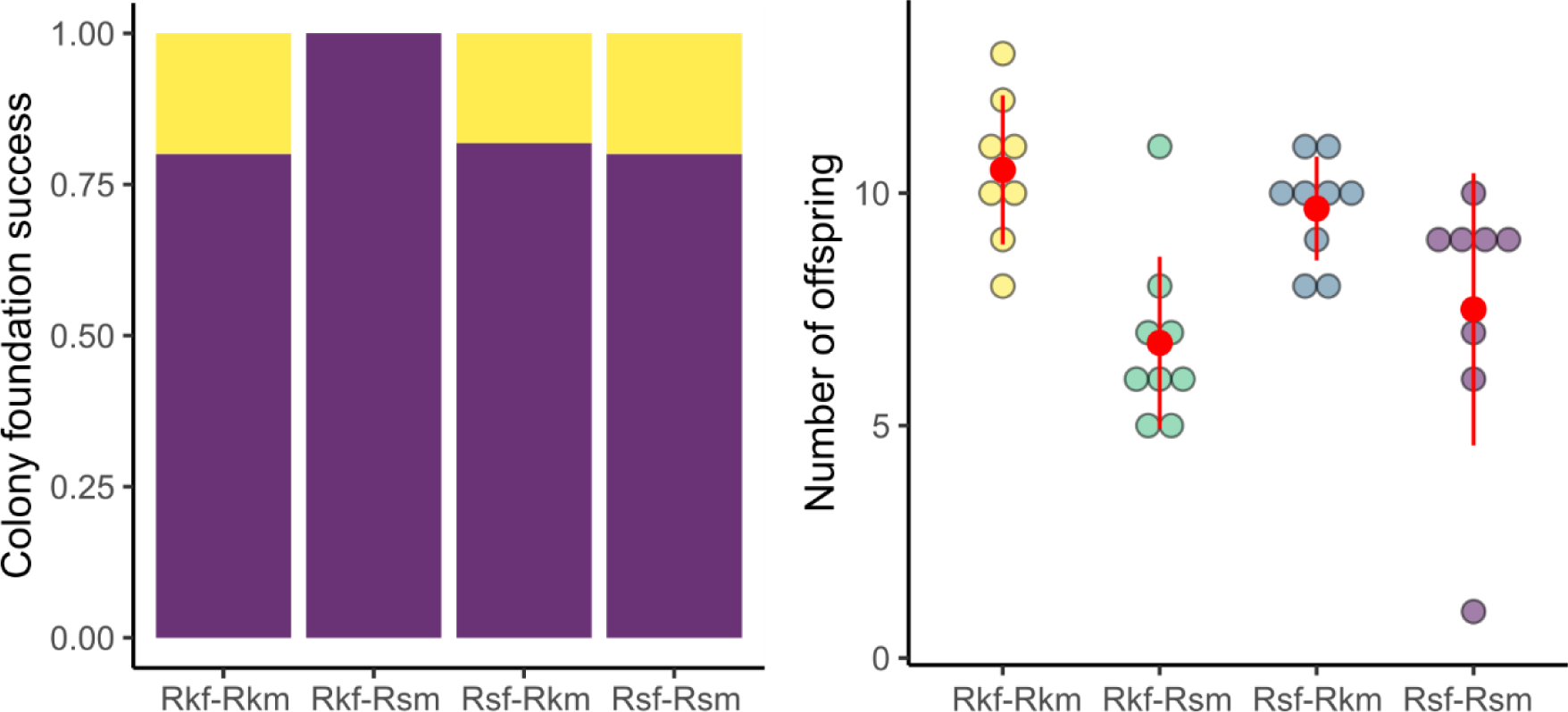
Comparison of colony foundation success in heterospecific pairs.

## Notes

https://github.com/nobuaki-mzmt/Ret-kanmonensis-swarming

### Competing Interest Statement

The authors have declared no competing interest.

## References

1. Abe T. 1987. Evolution of life types in termites In: Kawano S, Connell J, Hidaka T, editors. Evolution and Coadaptation in Biotic Communities. Tokyo: University of Tokyo Press. pp. 125–148.

2. Aguilera-Olivares D, Flores-Prado L, Véliz D, Niemeyer HM. 2015. Mechanisms of inbreeding avoidance in the one-piece drywood termite Neotermes chilensis. Insect Soc 62:237–245. doi:10.1007/s00040-015-0399-1

3. Aihetasham A, Akhtar MS. 2008. Swarming Behaviour and Alate Sex-Ratio of Heterotermes indicola (Wasmann) (Isoptera: Rhinotermitidae). Pakistan Journal of Zoology 40:75–82.

4. Camazine S, Deneubourg J-L, Franks NR, Sneyd J, Theraulaz G, Bonabeau E. 2001. Self-organization in Biological Systems. Princeton: NJ: Princeton University Press.

5. Chouvenc T, Scheffrahn RH, Mullins AJ, Su N-Y. 2017. Flight phenology of two *Coptotermes* species (Isoptera: Rhinotermitidae) in southeastern Florida. Journal of Economic Entomology 56:291–312. doi:10.1093/jee/tox136

6. Dornhaus A, Powell S, Bengston S. 2012. Group Size and Its Effects on Collective Organization. Annual Review of Entomology 57:123–141. doi:10.1146/annurev-ento-120710-100604

7. Endo J, Takanashi T, Mukai H, Numata H. 2019. Egg-Cracking Vibration as a Cue for Stink Bug Siblings to Synchronize Hatching. Current Biology 29:143–148.e2. doi:10.1016/j.cub.2018.11.024

8. Garnier S, Gautrais J, Theraulaz G. 2007. The biological principles of swarm intelligence. Swarm Intell 1:3–31. doi:10.1007/s11721-007-0004-y

9. Haverty MI, Howard RW. 1981. Production of soldiers and maintenance of soldier proportions by laboratory experimental groups ofReticulitermes flavipes (Kollar) andReticulitermes virginicus (Banks) (Isoptera:Rhinotermitidae). Ins Soc 28:32–39. doi:10.1007/BF02223620

10. Hölldobler B, Wilson EO. 1990. The Ants. Harvard University Press.

11. Husseneder C, Simms DM, Ring DR. 2006. Genetic diversity and genotypic differentiation between the sexes in swarm aggregations decrease inbreeding in the Formosan subterranean termite. Insect Soc 53:212–219. doi:10.1007/s00040-005-0860-7

12. Ishii Y, Hasegawa E. 2013. The mechanism underlying the regulation of work-related behaviors in the monomorphic ant, Myrmica kotokui. Journal of Ethology 31:61–69. doi:10.1007/s10164-012-0349-6

13. Jeanson R, Dussutour A, Fourcassié V. 2012. Key factors for the emergence of collective decision in invertebrates. Frontiers in Neuroscience 6:121. doi:10.3389/fnins.2012.00121

14. Khan Z, Zhang M, Meng YF, Zhao J, Kong XH, Su XH, Xing LX. 2019. Alates of the termite Reticulitermes flaviceps feed independently during their 5-month residency in the natal colony. Insect Soc 66:425–433. doi:10.1007/s00040-019-00698-9

15. Kitade O, Hayashi Y. 2002. Localized distribution of an alien termite Reticulitermes kanmonensis (Isoptera: Rhinotermitidae). Entomological Science 5:197–201.

16. Kusaka A, Matsuura K. 2017. Allee effect in termite colony formation: influence of alate density and flight timing on pairing success and survivorship. Insectes Sociaux 65:17–24. doi:10.1007/s00040-017-0580-9

17. Lee W, Choi DS, Ji JY, Kim N, Han JM, Park SH, Lee S, Seo MS, Hwang WJ, Forschler BT, Takematsu Y, Lee YH. 2015. A new record of Reticulitermes kanmonensis Takematsu, 1999 (Isoptera: Rhinotermitidae) from Korea. Journal of Asia-Pacific Entomology 18:351–359. doi:10.1016/j.aspen.2015.04.006

18. Lucena EF, Silva IS, Monteiro SRP, Moura FMS, Vasconcellos A. 2022. Accumulated precipitation and air density are linked to termite (Blattodea) flight synchronism in a Seasonally Dry Tropical Forest in north-eastern Brazil. Austral Entomology 61:78–85. doi:10.1111/aen.12577

19. Martius C. 2003. Rainfall and air humidity: non-linear relationships with termite swarming in Amazonia. Amazoniana 17:387–397.

20. Mitaka Y, Akino T, Matsuura K. 2023. Development of a standard medium for culturing the termite Reticulitermes speratus. Insect Soc 70:265–274. doi:10.1007/s00040-023-00907-6

21. Mitchell JD. 2008. Swarming flights of the fungus-growing termite, Macrotermes natalensis (Haviland) (Isoptera: Macrotermitinae), and the environmental factors affecting their timing and duration. African Entomology 16:143–152. doi:10.10520/EJC32788

22. Mizumoto N, Bourguignon T. 2022. Light alters activity but does not disturb tandem coordination of termite mating pairs. Ecological Entomology een.13209. doi:10.1111/een.13209

23. Mizumoto N, Bourguignon T. 2020. Modern termites inherited the potential of collective construction from their common ancestor. Ecology and Evolution 10:6775–6784. doi:10.1002/ece3.6381

24. Mizumoto N, Bourguignon T, Bailey NW. 2022a. Ancestral sex-role plasticity facilitates the evolution of same-sex sexual behavior. Proceedings of the National Academy of Sciences of the United States of America 119:e2212401119. doi:10.1073/pnas.2212401119

25. Mizumoto N, Bourguignon T, Kanao T. 2022b. Termite nest evolution fostered social parasitism by termitophilous rove beetles. Evolution 76:1064–1072. doi:10.1111/evo.14457

26. Moser JC, Reeve JD, Bento JMS, Lucia TMCD, Cameron RS, Heck NM. 2004. Eye size and behaviour of day- and night-flying leafcutting ant alates. J Zool Lond 264: 69-75.

27. Neoh K-B, Lee C-Y. 2009. Flight Activity of Two Sympatric Termite Species, Macrotermes gilvus and Macrotermes carbonarius (Termitidae: Macrotermitinae). Environmental Entomology 38:1697–1706. doi:10.1603/022.038.0623

28. Nishide Y, Tanaka S. 2016. Desert locust, Schistocerca gregaria, eggs hatch in synchrony in a mass but not when separated. Behav Ecol Sociobiol 70:1507–1515. doi:10.1007/s00265-016-2159-2

29. Noordijk J, Morssinkhof R, Boer P, Schaffers AP, Heijerman Th, Sýkora KV. 2008. How ants find each other; temporal and spatial patterns in nuptial flights. Insect Soc 55:266–273. doi:10.1007/s00040-008-1002-9

30. Nutting WL. 1979. Termite Flight Periods: Strategies for Predator Avoidance? Sociobiology.

31. Nutting WL. 1969. 8 Flight and colony foundation. In: Krishna K, Weesner FM, editors. Biology of Termites. New York: Academic Press. pp. 233–282. doi:10.1016/B978-0-12-395529-6.50012-X

32. Puckett RT, Espinoza EM, Gold RE. 2014. Alate Trap-Based Assessment of Formosan Subterranean Termite (Isoptera: Rhinotermitidae) Dispersal Flight Phenology Associated With an Urbanized Barrier Island Ecosystem. Environ Entomol 43:868–876. doi:10.1603/EN13223

33. R Core Team. 2023. R: A language and environment for statistical computing.

34. Sands WA. 1965. Alate development and colony foundation in five species of Trinervitermes (Isoptera, Nasutitermitinæ) in Nigeria, West Africa. Insectes Sociaux 12:117–130. doi:10.1007/BF02223758

35. Sen-Sarma PK. 1962. Some observations on swarming in nature and colony foundation under laboratory conditions in Odontotermes assmuthi (Holmgren) at Dera Dun (Isoptera: Termitidae). Beiträge zur Entomologie 12:292–297. doi:10.21248/contrib.entomo1.12.3-4.292-297

36. Sugio K, Miyaguni Y, Yoshimura T. 2020. Synchronization of alate emergence among colonies and dispersal strategy in the Ryukyu dry-wood termite Neotermes sugioi (Isoptera: Kalotermitidae). Insectes Sociaux 1–8. doi:10.1007/s00040-020-00766-5

37. Takematsu Y. 1999. The Genus Reticulitermes (Isoptera: Rhinotermitidae) in Japan, with description of a new species. Entomological Science 2:231–234.

38. Vergne AL, Mathevon N. 2008. Crocodile egg sounds signal hatching time. Current Biology 18:R513–R514. doi:10.1016/j.cub.2008.04.011

39. Vince MA. 1966. Artificial acceleration of hatching in quail embryos. Animal Behaviour 14:389–394. doi:10.1016/S0003-3472(66)80034-9

40. Wu C-C, Tsai C-L, Liang W-R, Takematsu Y, Li H-F. 2019. Identification of Subterranean Termite Genus, Reticulitermes (Blattodea: Rhinotermitidae) in Taiwan. Journal of Economic Entomology 112:2872–2881. doi:10.1093/jee/toz183

41. Wu J, Xu H, Hassan A, Huang Q. 2019. Interspecific Hybridization between the Two Sympatric Termite Reticulitermes Species under Laboratory Conditions. Insects 11:14. doi:10.3390/insects11010014

42. Wu Y, Chen J, Takata M, Matsuura K. 2023. Maternal determination of soldier proportion and paternal determination of soldier sex ratio in hybrid Reticulitermes (Isoptera: Rhinotermitidae) termite colonies. PLoS ONE 18:e0293096. doi:10.1371/journal.pone.0293096

